# How formate dehydrogenase-lacking acetogen *Clostridium bovifaecis* utilize formate as the sole carbon source for its acetogenic growth?

**DOI:** 10.1101/2023.06.12.544627

**Authors:** Ziwei Guan, Bo Fu, Ralf Conrad, Qingqing Qian, Dongfei Han, Hongbo Liu, He Liu

**Affiliations:** School of Environmental and Civil Engineering, Jiangsu Key Laboratory of Anaerobic Biotechnology, Jiangnan University, Wuxi, China; Jiangsu Collaborative Innovation Center of Technology and Material of Water Treatment, Suzhou University of Science and Technology, Suzhou, China; Max Planck Institute for Terrestrial Microbiology, Marburg, Germany

**Keywords:** Acetogen, Formate metabolism, Formate dehydrogenase, Ribonucleotide reductase, Wood-Ljungdahl pathway

## Abstract

Little is known about the growth of formate dehydrogenase-lacking acetogen on formate as sole carbon source. Here, we analyzed formate metabolism in *Clostridium bovifaecis* strain BXX using different concentrations of formate. The results show that *C. bovifaecis* converted formate (11.5-96 mM) into acetate with molar ratio of 2.0:1∼2.6:1 by using L-cysteine in the anaerobic medium as electron source according to the stoichiometry of acetogenesis. Genome analysis of *C. bovifaecis* revealed genes encoding anaerobic ribonucleoside triphosphate reductase (*nrdD* and *nrdG*) catalyzing the oxidation of formate to CO_2_ while ATP is being reduced to the desoxy form. The existence of *nrdD* was verified by PCR, reverse transcription-PCR analysis and acetogenesis from formate. The process mode of acetogenesis from formate in *C. bovifaecis* provides insight into the unique metabolic feature of an FDH-lacking acetogen.

**IMPORTANCE:** Wood-Ljungdahl pathway (WLP) lacking formate dehydrogenase (FDH) which catalyzes CO_2_ reduction to formate has been reported to occur acetogenesis only in the presence of formate and exogenous CO_2_, which seems to result from the formate-rich habitats adaptation of gastrointestinal acetogens. Here, we found FDH-lacking *Clostridium bovifaecis* strain BXX converted formate (11.5-96 mM) into acetate with molar ratio of 2.0:1∼2.6:1 fitting the stoichiometry of acetogenesis when using formate as the sole carbon source. CO_2_ needed in the carbonyl branch of WLP was from the oxidation of formate to CO_2_ catalyzed by anaerobic ribonucleoside triphosphate reductase while ATP is being reduced to the desoxy form. L-cysteine in the anaerobic medium was the electron source of WLP. The process mode of acetogenesis from formate in *C. bovifaecis* provides insight into how an FDH-lacking acetogen can make a living from the simplest resources as carbon source, which has both ecological and biotechnological significance.

## INTRODUCTION

Acetogenic bacteria are anaerobes using the Wood-Ljungdahl pathway (WLP) for both the dissimilation and assimilation of CO_2_ as well as using it as terminal electron acceptor for energy conservation [1]. Acetogens are phylogenetically diverse bacteria present in 23 different genera, but all have the common metabolic feature of converting 2 mol CO_2_ into 1 mol acetyl-CoA via WLP [2]. Our previous study showed that the fecal acetogen *Clostridium bovifaecis* strain BXX lacks a gene encoding formate dehydrogenase, which catalyzes the interconversion of CO_2_ and formate, and allows the acetogenic utilization of glucose only in the presence of formate and exogenous CO_2_ [3]. The substrate/product stoichiometry of formate/acetate was 2.75:1 being indicative of acetogenesis during growth on glucose-CO_2_-formate. The molar ratios of consumed formate and CO_2_ was consistent with a stoichiometry of 1:1 suggesting that formate instead of CO_2_ acted as electron acceptor for the FDH-lacking methyl branch of WLP. Although the previous study provided insight into the mechanism of formate utilization of the FDH-lacking acetogen using glucose-CO_2_-formate as mixed carbon sources, we are still curious about formate metabolism in strain BXX when using formate as the sole carbon source. However, little is known about formate metabolism in FDH-lacking acetogens. Also note that formate not only provides bacteria with a carbon source, but also represents an electron donor (reducing power) [4].

Formate can be produced from carbon dioxide and renewable energy sources [5]-[7], which is essential for establishing a sustainable carbon cycle economy [8]. As C1 compound, formate is easier to store and transport compared to C1 gases due to high water solubility, and high mass transfer. Thus, formate has received wide attention as a potential industrial feedstock for microbial growth [9],[10]. Studies on formate metabolism in the FDH-lacking acetogen is therefore of high scientific and biotechnological interest.

The model acetogen *Acetobacterium woodii* was recently reported to grow on 50-200 mM formate as a sole carbon source and to convert the formate to acetate at a ratio of 4.4:1, where 3 mol formate was oxidized to CO_2_ and provided electrons for the reduction of 1 mol formate and 1 mol CO_2_ to acetate [11]. Here, we studied acetogenesis from formate in FDH-lacking *C. bovifaecis*.

## MATERIALS AND METHODS

### 2.1 Strains and medium

*C. bovifaecis* strains BXX used in this study has been isolated and is stored by our laboratory [12]. The composition of basic medium was the same in the descriptions of Yao et al. [3] but without casitone, and the anaerobic medium contained 1.75 mM cysteine.

### 2.2 Growth of *C. bovifaecis* on formate-plus-CO_2_ or formate-only

Incubation experiments were performed in 120 mL serum bottles containing 50 mL of medium at 30°C, unless noted otherwise. For the growth on formate and formate-plus-CO_2_, the experiments consisted of two parallel incubations with different substrates as follows: (i) 46 mM formate, (ii) 46 mM formate and exogenous CO_2_. The exogenous CO_2_ was added by flushing with N_2_-plus-CO_2_ (80:20, v/v) gas to 10^5^ Pa overpressures. For the growth on formate as sole carbon source, parallel incubations with three concentrations of formate were as follows: (i) 11.5 mM formate, (ii) 69 mM formate, (iii) 138 mM formate. Bacterial growth was determined by measuring optical density at 600 nm (OD_600_). The experiments were carried out in triplicates.

### 2.3 Growth of *C. bovifaecis* on formate as the sole carbon source in the presence of different concentration of cysteine

To determine the electron source of *C. bovifaecis* when using formate as the sole carbon source, different concentrations of L-cysteine were added in the medium: (i) 0 mM, (ii) 11.5 mM, (iii) 23 mM. Bacterial growth was measured as described above. All experiments were performed in triplicates.

### 2.4 Chemical analysis

Liquid samples (3 mL) were collected at 0, 1, 3, 5 and 8 d. The concentration of formate was determined by using high-performance liquid chromatography (HPLC). An aliquot of 1 mL was centrifuged at 10,000×g for 5 min and then filtered through a 0.22 μm water system membrane. The mobile phase was a mixed solution of 0.01 mol/L KH_2_PO_4_ and methanol (95:5, v/v) with a flow rate of 1.0 mL/min. The detection wavelength, column temperature, injection volume was 206 nm, 30°C and 20 μl, respectively [3]. The concentration of acetate was determined by gas chromatography. The liquid samples were first mixed with an equal volume of 3 mM phosphoric acid and filtered through a 0.22-μm water system membrane. More details were described in Liu et al. [13]. CO_2_ concentrations were determined using gas chromatography (Foley, China) [14]. The SO_4_^2-^ concentration was determined using a barium sulfate turbidimetry as described in Wang et al.[15].

The concentrations of L-cysteine were measured with an ultraviolet-visible spectrophotometric assay [16]. Standard solutions of 0.2, 0.3, 0.4, 0.8 and 1.0 g/L of L-cysteine hydrochloride monohydrate were prepared. The composition of assay mixtures were (per 50 mL distilled water): 3.5 mL of NH_4_Fe(SO_4_)_2_ (0.02 M), 3.5 mL of orthophenanthroline (0.03 M), 1 mL of samples at different time points, 5 mL of triethanolamine (0.1 M)-HCl (1 M) buffer solution. The assay mixtures were incubated in a water bath at 25 °C for 50 min. After that, 1 mL of the above solution was transferred to a 25 mL volumetric flask, diluted with distilled water to volume. The absorbance was measured at 510 nm by UV spectrophotometer against a blank solution.

The concentrations of serine, glutamate, alanine, histidine and phenylalanine in the medium were determined with an HPLC system (Agilent 1100, Agilent Technologies, USA) [17]. An Agilent Hypersil ODS column (5 μm, 4.0 mm×250 mm) was used to separate the amino acids. The mobile phase A was a solution of sodium acetate (27.6 mM), trimethylamine and tetrahydrofuran (500:0.11:2.5, v/v, pH 7.2) and the mobile phase B was a solution of sodium acetate (0.9 mM), methanol and acetonitrile (1:2:2, v/v, pH 7.2) with a flow rate of 1.0 mL/min. The detection wavelength and column temperature were 338 nm and 40°C, respectively. Amino acid standards (l mM), O-phthalaldehyde (OPA) and 9-fluorenylmethyl chloroformate (FMOC) were purchased from Sigma Company. Liquid samples were mixed with an equal volume of trichloroacetic acid (TCA, 5%(v/v)) and filtered through a 0.22-μm water system membrane before the determination.

### 2.5 RNA extraction and PCR

Cells of *C. bovifaecis* in the early exponential growth phase were harvested for RNA extraction. Total RNA was extracted with an RNAprep pure cell/bacteria Kit. Reverse transcription used PrimeScript RT reagent kit with genomic DNA (gDNA) Eraser. The concentration of RNA and cDNA were determined by micro-spectrophotometer (Thermo NanoDrop ND 2000, USA). Anaerobic ribonucleoside triphosphate reductase expression levels were quantified by amplification of *nrdD* gene using the primers of *nrdD*-f (5’-ACAACTGGCTCGTGCTTACA-3’) and *nrdD*-r (5’-GTGCATCCGGGCTACCTAAA-3’). All oligonucleotide primers were synthesized and kits were purchased from Shanghai Sangon Biotech Co., Ltd. (China). The reverse transcription quantitative PCR (RT-qPCR) conditions for the *nrdD* gene were initialized at 94°C for 5 min, followed by 35 cycles at 94°C for 30 s, 55°C for 40 s and 72°C for 90 s [3]. Genomic DNA from *Acetobacterium woodii* was used as negative control of the *nrdD-f* and *nrdD-r* primers. Double-distilled water (ddH_2_O) was used as blank control.

## RESULTS

### 3.1 Growth of *C. bovifaecis* on formate-only or on formate-plus-CO_2_

To understand the acetogenic growth of *C. bovifaecis* we studied the growth on formate-only and on formate in the presence of CO_2_ (46 mM) (**Fig. 1**). In the formate-only incubations, about 4.0 mM of formate was consumed, accompanied by an acetate production of 1.56 mM, and the cell densities increased significantly with the final OD_600_ of approximately 0.10. In the formate-plus-CO_2_ incubations, acetate concentrations accumulated to a final concentration of 1.05 mM with formate consumption of 2.83 mM, but the OD_600_ did not increase (**Fig.1**), indicating that the bacterial growth was not active under this condition. No CO_2_ gas was detected in the headspace of the formate-only incubations, but small amounts of CO_2_ (0.70 mM) were consumed in the formate-plus-CO_2_ incubations.

**Fig. 1.**
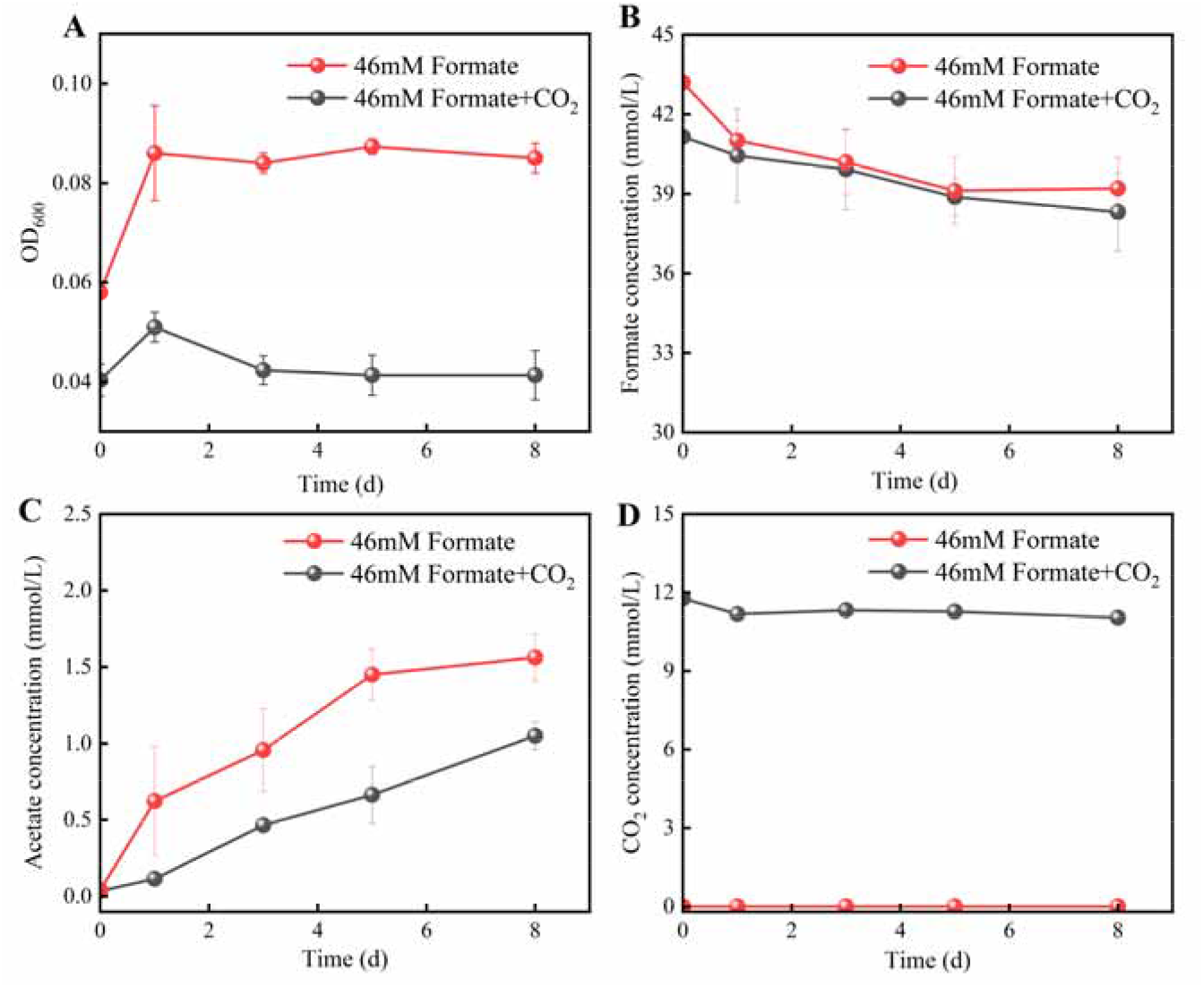
Cell density (A), formate consumption (B), acetate production (C) and CO_2_ concentration (D) of C. *bovifaecis* using 46 mM formate or with supplemental CO_2_ as substrates.

### 3.2 Growth of *C. bovifaecis* on different concentrations of formate

To understand the growth of *C. bovifaecis* on formate as sole carbon source, different concentrations of formate (11.5-138 mM) were used (**Fig. 2**). The amounts of formate utilization decreased with increasing formate concentration. When 11.5 mM and 69 mM formate was supplied, 5.43 mM and 2.95 mM formate was consumed with an acetate production of 2.25 mM and 1.21 mM, respectively (**Fig. 2**). The cell densities increased significantly with the final OD_600_ reaching approximately 0.10 to 0.15. However, no formate consumption was detected in the 138 mM formate incubations and the OD_600_ values did not increase (**Fig. 2)**. No CO_2_ gas was detected in all the incubations. The pH of all the incubations gradually decreased from approximately pH 7.00 to pH 6.50 (**Fig. 2**).

**Fig. 2.**
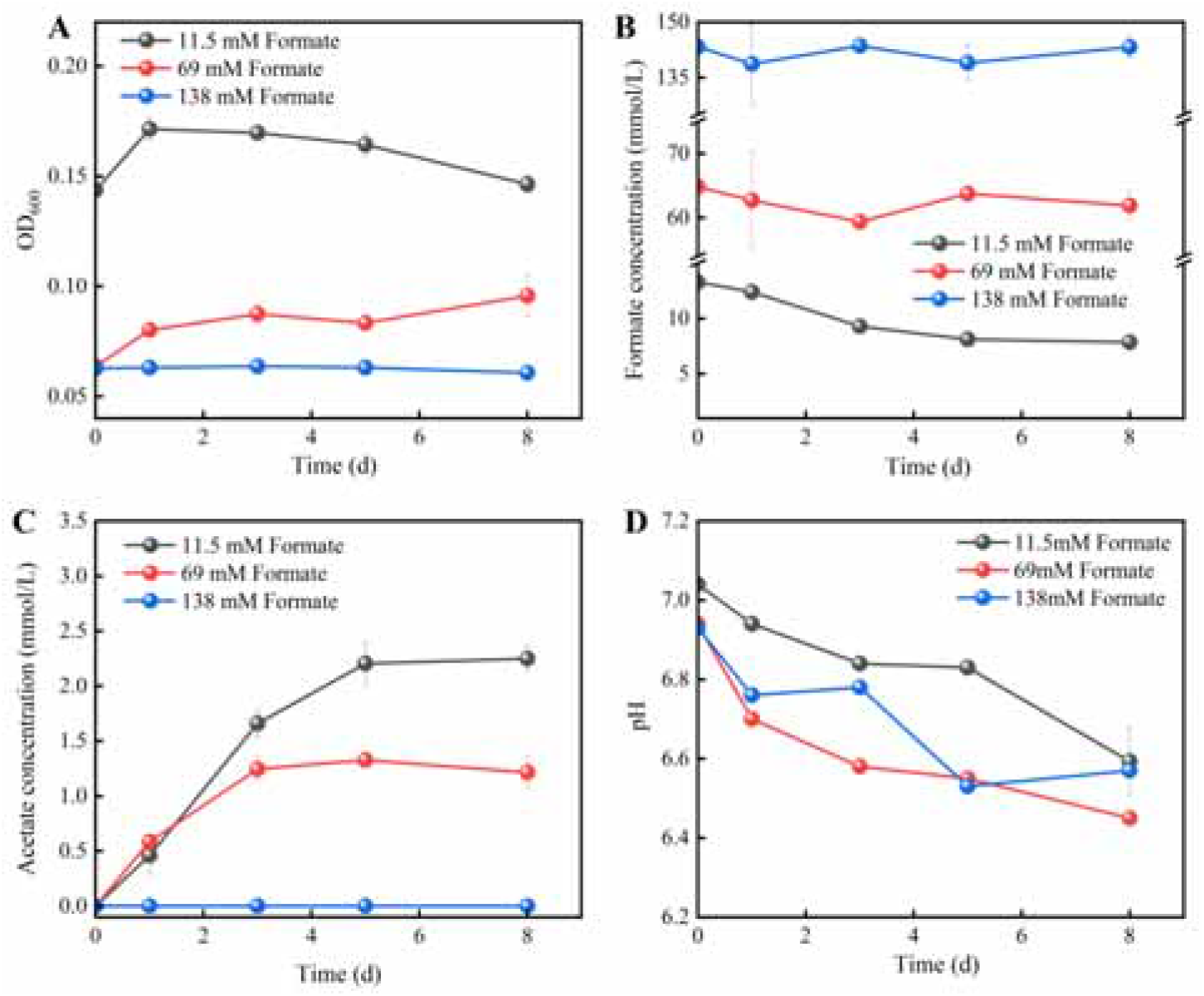
Cell density (A), formate consumption (B), acetate production (C) and pH value (D) of *C. bovifaecis* using 11.5 mM, 69 mM and 138 mM formate as the sole carbon source.

The amounts of consumed formate and of produced acetate and CO_2_ are summarized in **Table 1**. 47.2% of the initial formate was utilized when the initial formate was 11.5 mM, but less than 10% of the initial formate was consumed when formate was 46-69 mM. Formate was converted to acetate with a ratio of 2.4:1∼2.5:1 when the formate concentrations was 11.5-69 mM, which is consistent with the stoichiometry of WLP converting 1 mol formate and 1 mol CO_2_ into 1 mol acetate. In the presence of exogenous CO_2_, a slightly higher ratio of 2.8:1 was obtained.

**Table 1.**
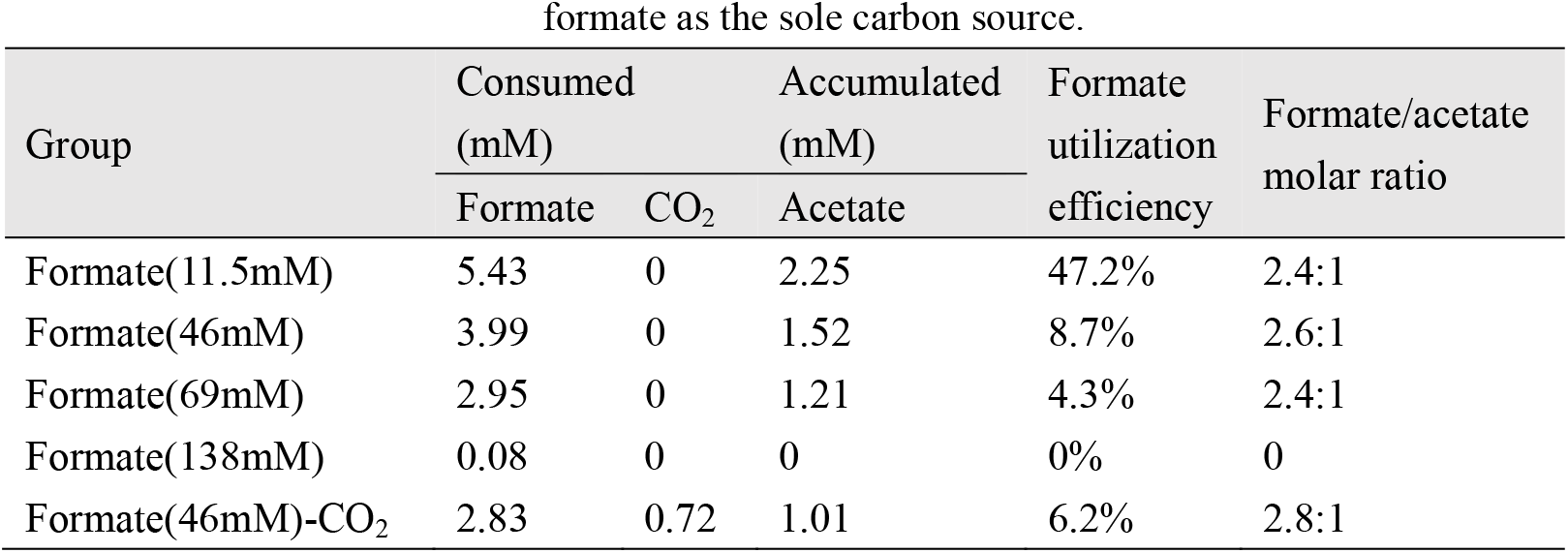
Accumulated or consumed metabolites in *C. bovifaecis* using different concentrations of formate as the sole carbon source.

### 3.3 Growth of *C. bovifaecis* on formate with different concentrations of cysteine

Cysteine is commonly used as a reducing agent in anaerobic medium [18], [19], and may be a potential electron source for *C. bovifaecis* using formate as carbon source. Here, different concentrations of L-cysteine (0-23 mM) were applied when formate was 46 mM (**Fig. 3**). The cell density increased significantly with the increase of cysteine concentration. The OD_600_ increased by 0.06 and 0.07 for 11.5 mM and 23 mM cysteine, respectively, while the OD_600_ remained unchanged when cysteine was absent (**Fig. 3A**). The pH of all the incubations was ranged from 7.0 to 8.70 (**Fig. 3B**). The consumption of formate, cysteine, and the production of acetate at different initial cysteine concentrations is shown in **Fig. 3C, 3D**, and **3E**, respectively. Cysteine of 1.75-6.42 mM were consumed when cysteine concentrations of 1.75 mM to 23 mM were supplied.

**Fig. 3.**
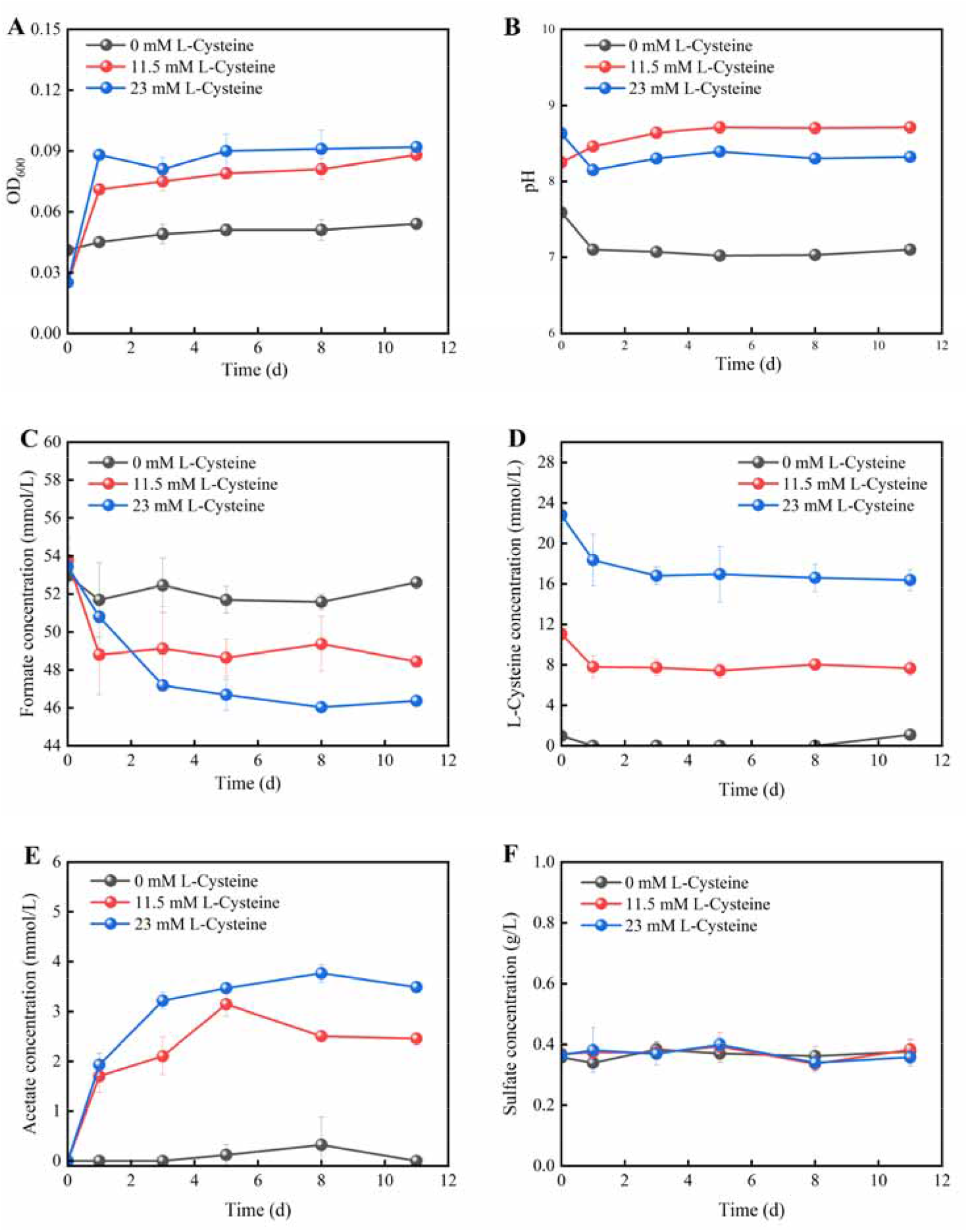
Cell density (A), pH value (B), formate consumption (C), cysteine consumption (D), acetate production (E) and sulfate concentration (F) of *C. bovifaecis* in incubations with formate as sole carbon source in presence of 0, 11.5 mM and 23 mM cysteine.

The balance of formate and cysteine consumption and acetate production are shown in **Table 2**. Formate consumption and acetate production increased with increasing cysteine concentrations, and the molar ratio of formate/acetate decreased from 2.6:1 to 2.0:1. Also the consumption of cysteine increased, and the molar ratio of cysteine/acetate increased from 1.1:1 to 1.8:1. No acetate was formed when cysteine was absent. No CO_2_ gas and sulfate production were detected in all the incubations.

**Table 2.** Accumulated or consumed metabolites in *C. bovifaecis* using 46 mM of formate as the sole carbon source in presence of different concentrations of cysteine.

| Group | Consumed<br>(mM) |  | Accumulated<br>(mM) | Formate<br>utilization<br>efficiency | Formate<br>/acetate | Cysteine<br>/acetate |
| --- | --- | --- | --- | --- | --- | --- |
|  | Formate | Cysteine | Acetate |  | molar<br>ratio | molar<br>ratio |
| Cysteine (0 mM) | 0.34 | 0 | 0 | 0.7% | 0 | 0 |
| Cysteine (1.75 mM) | 3.99 | 1.75 | 1.52 | 8.7% | 2.6:1 | 1.1:1 |
| Cysteine (11.5 mM) | 5.33 | 3.38 | 2.45 | 11.6% | 2.2:1 | 1.4:1 |
| Cysteine (23 mM) | 7.07 | 6.42 | 3.49 | 15.4% | 2.0:1 | 1.8:1 |

The sulfate concentration stayed constant at approximately 4 mM (**Fig. 3F**).

To further study the conversion of cysteine, the change of amino acids concentration in the medium was examined. Without addition of cysteine, only a trace amount of glutamate (0.08 mM) was detected. In the presence of 11.5 mM cysteine, an obvious increase of the serine concentration (0.30 mM) was observed. In presence of 11.5 mM or 23 mM cysteine, an obvious increase of the alanine and glutamate concentrations (0.22-0.35 mM) was observed, and small amounts of histidine and phenylalanine were also generated (**Table 3**).

**Table 3.** Accumulated amino acid in the medium with different concentrations of cysteine.

| Amino acid | Cysteine (0 mM) |  | Cysteine (11.5 mM) |  | Cysteine (23 mM) |  |
| --- | --- | --- | --- | --- | --- | --- |
|  | initial | end | initial | end | initial | end |
| Alanine | 0.02 | 0.01 | 0.02 | 0.25 | 0.02 | 0.24 |
| Glutamate | 0.18 | 0.27 | 0.19 | 0.54 | 0.19 | 0.43 |
| Serine | 0.01 | 0.01 | 0.02 | 0.30 | 0.02 | 0.01 |
| Histidine | 0.02 | 0.07 | 0.03 | 0.02 | 0.04 | 0.22 |
| Phenylalanine | 0.05 | 0.09 | 0.06 | 0.06 | 0.07 | 0.17 |

### 3.4 Detection of the *nrdD* gene on DNA and RNA level

*C. bovifaecis* is unable to oxidize formate to CO_2_ due to the lack of formate dehydrogenase. Anaerobic ribonucleoside-triphosphate reductase is a potential enzyme catalyzing the oxidation of formate to CO_2_ in *C. bovifaecis* strain BXX. Ribonucleotide reductase (RNR) enzymes catalyze the reduction of ribonucleotides to deoxyribonucleotides that are needed for DNA synthesis and DNA repair in all living cells [20], [21]. RNRs have been divided into three classes (I, II and III) based on the metallocofactors required to generate this initiating thiyl radical [22]. The class III

RNR enzymes use formate besides or instead reduced dithiols to provide the necessary electrons. Class III RNR enzymes are found in facultatively anaerobic bacteria, and expressed only under anaerobic conditions. Class III enzymes consist of two proteins, n*rdD* encoding for reductase and *nrdG* encoding for activating protein [23]. As expected, the genome of *C. bovifaecis* strain BXX contained both *nrdD* and *nrdG* genes.

To verify the presence and expression of the genes in *C. bovifaecis* strain BXX, the genes and transcripts of *nrdD* were amplified using PCR and reverse transcription PCR (RT-PCR) (**Fig. 3A**) and quantified using reverse transcription quantitative PCR (RT-qPCR) (**Fig. 3B**). As expected, the *nrdD* gene in *C. bovifaecis* strain BXX was existent and expressed during growth on 11.5 mM formate. The transcription copies of *nrdD* gene in *C. bovifaecis* increased by 1 magnitude during the initial 3 days, then decreased by 2 magnitudes to a final level of 10^4^ copies/mL.

## DISCUSSION

In the metabolic pathway of acetogenesis from formate in the model acetogen *Acetobacterium woodii*, 3 mol formate are oxidized to 3 mol CO_2_ and 3 mol H_2_ by a hydrogen-dependent carbon dioxide reductase (HDCR) containing a formate dehydrogenase (FDH) and a hydrogenase, and another 1 mol formate and 1 mol CO_2_ is reduced to generate acetyl-CoA via WLP, which is further converted to acetate [11].

*A. woodii* can convert 4.4 mol formate to 1 mol acetate [11]. In comparison, the molar ratio of formate consumed and acetate produced in *C. bovifaecis* strain BXX was 2.0:1∼2.5:1 (**Table 1**), close to a stoichiometry of 2:1, which indicates acetogenesis from 1 mol formate and 1 mol CO_2_. Formate dehydrogenase plays an important role in formate metabolism of *A. woodii*, whereas *C. bovifaecis* strain BXX lacks FDH. Anaerobic class III RNRs catalyzing the reduction of oxyribonucleoside triphosphates with formate as the electron donor provide *C. bovifaecis* with deoxyribonucleoside triphosphates required for DNA synthesis. *C. bovifaecis* converted 1 mol formate into 1 mol CO_2_ by ribonucleoside-triphosphate reductase (Eq.1) for ribonucleotide synthesis.

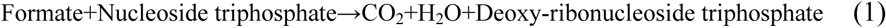

As DNA replication primarily occurs during cell division, cell growth with cellular dry weight of 0-0.0058 g·L^-1^ was relatively low with formate (11.5-138mM) as sole carbon source (**Table 4**). Similarly, this also may be the reason for the low formate utilization efficiency (0-8.7%) under higher formate concentration (69-138 mM). Generally, the bioconversions are a mixture of catabolism and anabolism, which usually have completely different quantities. But in fact, catabolism and anabolism of *C. bovifaecis* in similar proportions using formate as sole carbon source may be due to the low molar growth yield of 0-5.89 g dw (mol formate)^-1^ (**Table 4**). By contrast, the acetogen *Peptostreptococcus productus* grew on fructose with a molar growth yield of 36-62 g dw (mol fructose)^-1^ [24].

**Table 4.** Growth yield of *C. bovifaecis* with different concentrations of formate

| Group | OD<sub>600</sub> | Dry weight (g·L<sup>-1</sup>) | Formate consumed (mM) | Molar growth yield<sup>a</sup> (g·mol<sup>-1</sup>) |
| --- | --- | --- | --- | --- |
| Formate(11.5mM)-Cysteine (1.75 mM) | 0.028 | 0.005 | 5.43 | 0.92 |
| Formate(46mM)-Cysteine (1.75 mM) | 0.027 | 0.005 | 3.99 | 1.20 |
| Formate(69mM)-Cysteine (1.75 mM) | 0.032 | 0.006 | 2.95 | 1.97 |
| Formate(138mM)-Cysteine (1.75 mM) | 0 | 0 | 0.08 | 0 |
| Formate(46mM)-CO<sub>2</sub>-Cysteine (1.75 mM) | 0.001 | 0 | 2.83 | 0 |
| Formate(46mM)-Cysteine (0 mM) | 0.013 | 0.002 | 0.34 | 5.89 |
| Formate(46mM)-Cysteine (11.5 mM) | 0.063 | 0.012 | 5.33 | 2.25 |
| Formate(46mM)-Cysteine (23 mM) | 0.067 | 0.013 | 7.07 | 1.81 |

<sup>a</sup> Expressed in grams (dry weight) of cells mole of substrate consumed<sup>-1</sup>.

Besides the function of formate being oxidized to CO_2_, the other function involves the condensation of formate with another metabolic intermediate [25]. Formate-tetrahydrofolate (THF) ligase is the only known formate-fixing reaction supporting the growth on formate as the sole carbon source. In the WLP of acetogen, formyl-THF is reduced to methyl-THF via the intermediate methylene-THF. Methyl-THF reacts with CO_2_ and CoA, resulting in the production of acetyl-CoA, which is further converted to acetate (acetogenesis) and assimilated into central metabolism. However, this conversion requires electron equivalents, which apparently were not derived from formate. Therefore, an alternative source for electron equivalents is required.

Göbbels et al. demonstrated that cysteine can be used as an electron source that contributes to acetate production in the CdS-*M. thermoacetica* biohybrid system [19]. Cysteine in the medium can be an electron source for *C. bovifaecis* with formate as the sole carbon source. In our study, there was no acetate production in formate-only incubations without cysteine, whereas formate was converted to acetate in the presence of cysteine. The ratio of formate/acetate increased from 2.6:1 to 2.0:1 with the increase of cysteine concentration from 1.75 to 23 mM. Previous studies have shown that some microorganisms can use cysteine as a carbon source in addition to using it as an energy source forming acetate from cysteine [19], [26]. Theoretically, if cysteine is used as a carbon source in our work, 1 mol cysteine should be converted to 1 mol acetate [27], and then formate should be converted to acetate with a ratio of 1:1. However, although increasing cysteine concentrations improved the formate utilization of *C. bovifaecis*, it did not result in higher acetate production. In addition, only 0.32 and 0.50 mM of acetate was produced for the incubations with 23 mM cysteine or alanine in absence of formate, respectively (data not shown). It is shown that acetate was mainly generated from formate but not cysteine, which only provided electrons for formate metabolism via the WLP. In other words, acetogenic growth of FDH-lacking *C. bovifaecis* occurred with formate as the carbon source and cysteine as the electron source. Based on the elemental balance, the whole metabolic reaction might be according to the following equation (Eq. 2):

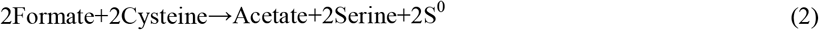

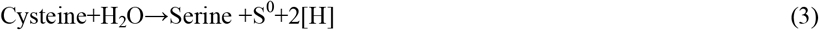

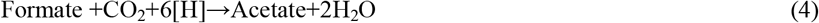

As cysteine is a core compound in many metabolic processes such as sulfur metabolism and protein biosynthesis, it can be converted to other amino acids by different enzymes [19]. Combining metabolic data and gene annotations of the genome in *C. bovifaecis*, the possible pathway of electron generation is proposed as follows: *C. bovifaecis* converted 1 mol L-cysteine to L-serine and sulfane sulfur, and produced reducing equivalents providing the electrons needed for WLP (Eq. 3). Subsequently, 1 mol formate and 1 mol CO_2_ was converted to acetyl-CoA by WLP, finally generating acetate (Eq. 4). Consistently, a utilization of cysteine was accompanied by an increase of the serine concentration. However, the increase of serine in the medium was lower than the assumed stoichiometry of 1:1 for produced serine and consumed cysteine. The produced serine may be used for protein synthesis and cell growth [28], so that only small amounts of serine were detected in the medium with 11.5 mM cysteine. The fate of S^0^ may be similar to that of serine, and the S^0^ may form intracellular inclusion such as sulfur granules stored in the cell [28], [30]. Given formate as the sole carbon source may cause nutritional limitation for the FDH-lacking acetogen, the formation of inclusions can be used as carbon or energy source for the future existence.

Overall, the results suggested the model for acetogenesis from formate in FDH-lacking acetogen *C. bovifaecis* as shown in **Fig. 5**. 1 mol formate is oxidized to 1 mol CO_2_ by anaerobic ribonucleoside triphosphate reductase, another 1 mol formate is converted to formyl-THF via formyl-THF synthase. Formyl-THF is further reduced to methyl-THF. 1 mol CO_2_ is reduced to CO and combined with the methyl-THF group by the CODH/ACS complex, generating acetyl-CoA. Acetyl-CoA is further converted to acetate. 2 mol L-cysteine was converted into serine and sulfane sulfur, and provided the electrons needed for WLP.

**Fig. 4.**
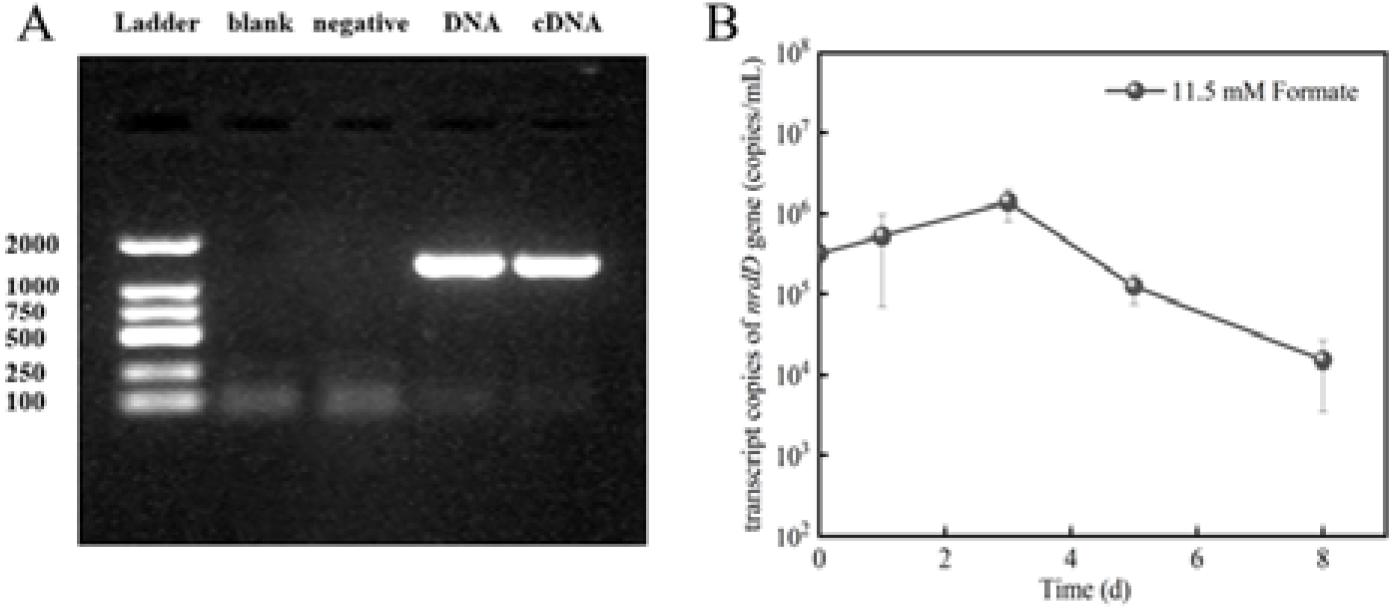
Gel electrophoresis images of PCR and RT-PCR products (A) and transcription copies (B) of *nrdD* gene during growth on 11.5 mM formate by *C. bovifaecis*. Genomic DNA from *Acetobacterium woodii* was used as negative controls and double-distilled water (dd H_2_O) was used as blank.

**Fig. 5.**
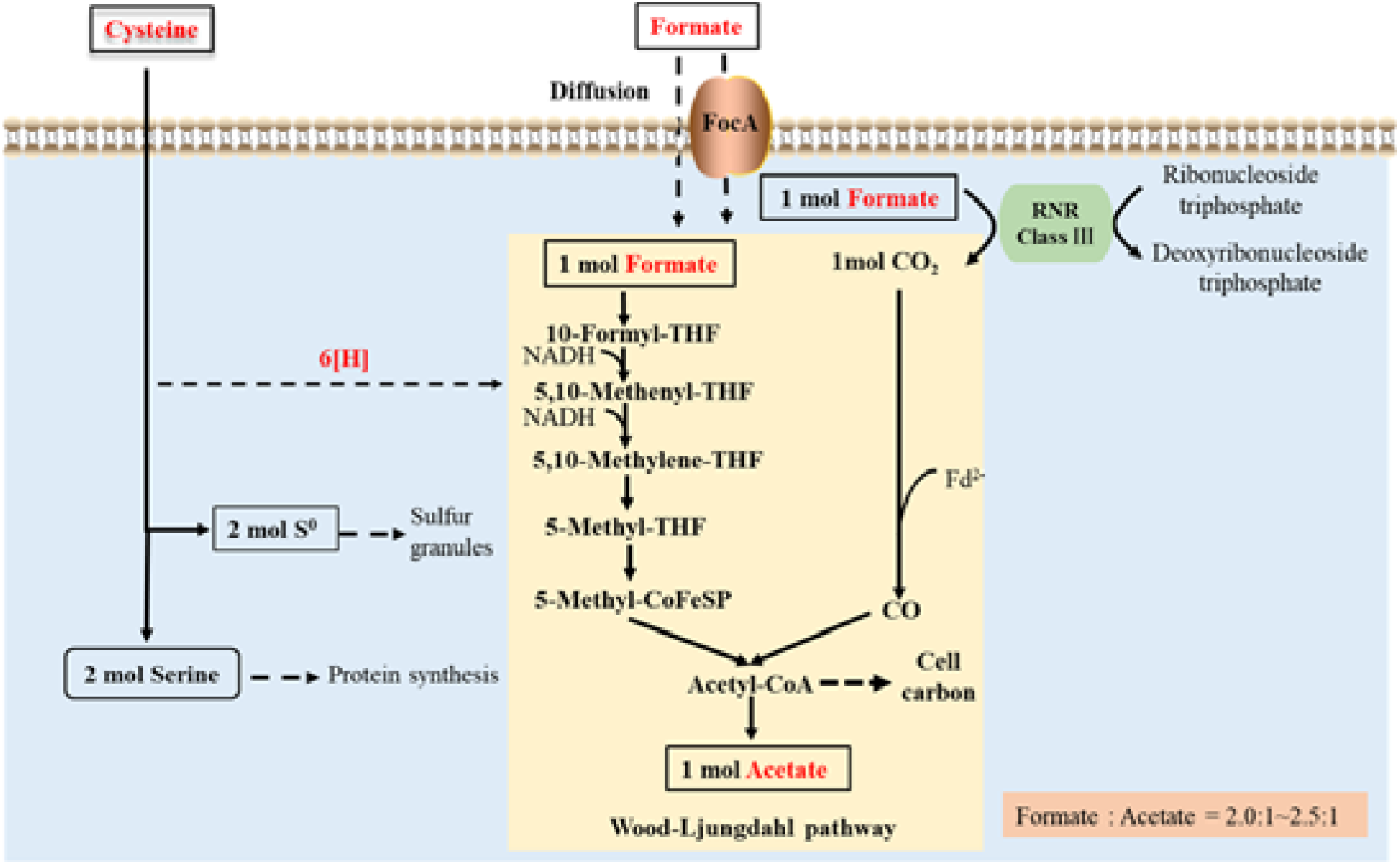
Acetogenesis from formate in FDH-lacking *C. bovifaecis*. Formate is imported via passive diffusion or the formate channel A (FocA). 1 mol formate are oxidized to 1 mol CO_2_ by anaerobic ribonucleoside-triphosphate reductase (RNR Class ?). 1 mol formate and 1 mol CO_2_ generate acetyl-CoA via WLP, which is further converted to 1 mol acetate. L-cysteine is converted into 2 mol sulfane sulfur which provided the electrons for WLP during acetogenesis. Sulfane sulfur further forms intracellular inclusion such as sulfur granules. The produced serine may be used for the protein biosynthesis.

Surprisingly, 138 mM formate did not support cell growth (**Fig. 2A**) and no acetate was produced (**Fig. 2C**). The reason may be that too high concentration of formate leads to severe acidification of the cell cytoplasm and consequent cell damage [31]. The uptake is the first step in formate metabolism. The pH of all the incubations gradually decreased from 7.0 to 6.50 (**Fig. 2D**). At physiological pH, formate is mainly present in the dissociated species and therefore it cannot diffuse across the cytoplasmic membrane but requires a transport system. One gene encoding for formate-nitrite transporter channel, *focA*, was annotated in the genome of *C. bovifaecis*. For active transport of formate, *C. bovifaecis* is like *E. coli* in that formate enters the cell via a putative channel *focA* [32].

Formate metabolism in *C. bovifaecis* is a study of how an FDH-lacking acetogen can make a living from the simplest resources as carbon source, which has both ecological and biotechnological significance. In terms of microbial ecology, formate is a common fermentation product of carbohydrate, which tends to accumulate in anaerobic environments lacking a terminal electron acceptor. Particularly in primordial environment with high concentration of formate under alkaline conditions [33], [34], the acetogenic growth of FDH-lacking acetogen on formate as the sole carbon source allow the conservation of energy supporting early life [35], and generate acetate served as a substrate for downstream metabolic processes. The role of cysteine as an electron source was emphasized in this study. Studies have shown the presence of cysteine-synthesizing microorganisms in the rumen [36], which may provide electron source for *C. bovifaecis* with formate as a carbon source. Cysteine has the potential to prevent nitrite poisoning in ruminant, which is related to the role of cysteine as an electron source for microbial metabolism to reducing nitrate. In terms of biotechnological application, the high conversion rate of formate into acetate in *C. bovifaecis* provide a potential advantage for microbial production of value-added chemicals from liquid or wastewater with low concentration of formate or formate-plus-CO [37], as formate can be produced efficiently from multiple available sources (e.g. electricity and biomass) and CO are the main constituent of synthesis gas.

## ACKNOWLEDGEMENTS

This research is supported by the Natural Science Foundation of Jiangsu Province (BK20181344) and the National Natural Science Foundation of China (No. 51978313). The authors gratefully acknowledge Mengxuan Su and Yushuai Cheng for their helpful experiment assistance.

